# Host competence–abundance relationships drive the dilution effect across multiple small mammal-borne pathogens

**DOI:** 10.1101/2025.11.28.691242

**Authors:** Marie Bouilloud, Maxime Galan, Julien Pradel, Vincent Sluydts, Nathalie Charbonnel, Benjamin Roche

## Abstract

The "dilution effect" hypothesis suggests that high host diversity can reduce the prevalence of certain zoonotic pathogens. Despite growing interest, its generality and practical relevance remain debated. In particular most previous studies have focused on large scale analyses or meta-analyses, limiting their potential use for zoonotic outbreak prevention. Here, we present an extensive field-based study of small mammals and their multiple pathogens to assess diversity-disease relationships at a local scale, revealing pathogen-specific patterns. While no significant association was detected between the prevalence of some pathogens and host diversity, others such as *Orthopoxvirus* and a strain of *Mycoplasma haemomuris* exhibited dilution effect patterns, whereas another *Mycoplasma haemomuris* strain and *Mycoplasma coccoides* displayed amplification effects. Crucially, we demonstrate that these contrasting outcomes are consistently explained by the host competence-abundance relationship. Our findings underscore the importance of host community composition and highlight the need to consider multiple pathogens when evaluating the ecological mechanisms underpinning the diversity-disease relationships.

## Introduction

The concomitant acceleration of infectious disease emergence 1 and biodiversity decline due to anthropogenic changes 2 suggests a strong connection between environmental conditions, host communities structure and pathogen circulation 3. Multiple meta-analyses emphasize that increased host diversity can reduce pathogen prevalence across a variety of host-pathogen systems 4–6. This supports the ‘dilution effect’ hypothesis, which posits that diversity can mitigate the transmission of zoonotic pathogens 7. Consequently, emerging research suggest that managing diversity, including species control 8 and conservation measures 9, may establish a favorable balance between diversity and public health 10.

However, these meta-analyses have been criticized for their reliance on non-uniform comparisons across studies, which can introduce biases and lead to misinterpretations related to differences in spatial scale, methods and metrics used 11–14. Furthermore, some studies have shown that the relationship between diversity and pathogen prevalence could be also neutral or even positive (amplification effect) 7,15, depending strongly on the local ecological context 11,16. Therefore, implementing biodiversity-based management strategies without accounting for system-specific features could lead to unwilled and potentially detrimental outcomes, particularly in ecosystems where the dilution effect does not apply. To address this uncertainty, there is crucial need for empirical studies that assess the prevalence of multiple pathogens across host species within a unified experimental framework. Such approaches are essential to identify the ecological conditions under which pathogen prevalence increases or decreases, thereby informing win-win strategies that simultaneously support both biodiversity conservation and public health objectives.

The theoretical mechanisms underlying the dilution effect are well described in the literature 7,17,18. First, host species within a community must vary in their competence for a given pathogen, i.e., their ability to get infected, to replicate and then disseminate the pathogen 19. Second, diversity must regulate the frequency of host encountering events, thereby reducing the transmission rate for competent host species and/or the abundance of competent hosts via ecological interactions 7,17,20. Third, the less abundant species tend to be lost first in declining biodiversity contexts, leaving communities dominated by more competent hosts 21,22 Finally, the pathogen must be horizontally transmitted, as vertically transmitted pathogens are less influenced by host diversity 23. These conditions are not mandatory, but they increase the likelihood of observing a dilution effect. To identify local contexts where the dilution effect may occur, it is crucial to empirically evaluate the relative contribution of each of these factors 24. This need is particularly critical given that host species competence remains understudied in empirical research, likely due to the methodological challenges associated with its measurement 25.

In this study, we investigated the local complex interactions among small mammal community diversity, host competence, and pathogen prevalence along a gradient of forest anthropization for multiple small mammal borne pathogens. Our primary objective was to determine whether a dilution effect might occur in these communities across most pathogens, or if diversity-prevalence relationships were pathogen specific. To address this question, we conducted a single, comprehensive empirical study examining the correlation between small mammal species diversity and the prevalence of seven small mammal pathogens across multiple forested sites. In a second step, we explored the ecological mechanisms potentially driving the patterns observed. Specifically, we hypothesized that host species would differ in their competence for each pathogen, as that dilution effects would emerge when competent hosts were more abundant in low diversity communities. To test this, we developed a quantitative host competence index and assessed the relationship between species competence, abundance, and pathogen prevalence.

## Results

### A large dataset of small mammals and their pathogens across forested sites

To capture a broad range of host species diversity, we conducted field sampling across five distinct periods, three during the breeding season and two during the post-breeding season, at six forested sites differing in their geography, climate, and levels of anthropization (Supplementary note 1, Fig. S1). Overall, we sampled 23 small mammal communities, including 16 species from different taxonomic groups. Among these, 11 species belonged to the order *Rodentia* with *Glis glis, Sciurus vulgaris, Rattus norvegicus*, *Mus musculus*, *Apodemus sylvaticus*, and *Apodemus flavicollis*, *Clethrionomys* (*syn. Myodes*) *glareolus, Microtus arvalis, Microtus subterraneus*, and *Microtus agrestis*. Additionally, six insectivore species were identified, *Sorex araneus*, *Sorex coronatus*, *Neomys fodiens*, *Crocidura russula*, and *Crocidura leucodon*, and *Erinaceus europaeus*. Species composition varied with the level of anthropization, while species richness, abundance and community diversity fluctuated primarily across sampling periods (Supplementary note 1, Figs. S2 and S3).

We identified sixteen pathogens from 1,549 small mammals examined using a combination of molecular and serological methods: 16S RNA V4 region metabarcoding for pathogenic bacteria, qPCR targeting the lip32 gene for pathogenic *Leptospira* sp., and immunofluorescence assays for detection of antibodies against viruses^26^. These pathogens varied in their taxonomic classification, transmission mode (direct vs vector-borne), and host specificity. Pathogen distribution varied across sites. Only seven of them were present at all sites. These included three strains of *Mycoplasma haemomuris* (designated *Mhae1, Mhae2, Mhae3,* Fig. S4), one strain of *Mycoplasma coccoïdes (Mcoc)*, and pathogenic species of *Leptospirosa (Lept), Bartonella (Bart)* and an orthopoxvirus (POXV). *Leptospira* and POXV are directly transmitted pathogens, while the others are vector-transmitted. For the rest of this study, we focused our analyses on these seven widely distributed pathogens.

### Contrasted relationships between pathogens presence and small mammal diversity

We assessed the relationship between diversity (quantified through the Shannon index) and the prevalence of each pathogen, while controlling for confounding factors including environmental variables, host sex and functional age. Our analyses revealed that the relationship between pathogen prevalence and small mammal diversity was pathogen specific, with the direction and strength of the relationship varying across the different pathogens considered.

We observed a negative correlation between the presence of anti-POXV antibodies (Wald test, β = -2.87, z = 6.59, p =< 2e-16) as well as *M. haemomuris* strain 2 (Wald test, β = -1.24, z = -3.53, p = 4.00e-04) and small mammal diversity (Fig. 1, Table S1).

**Fig. 1.**
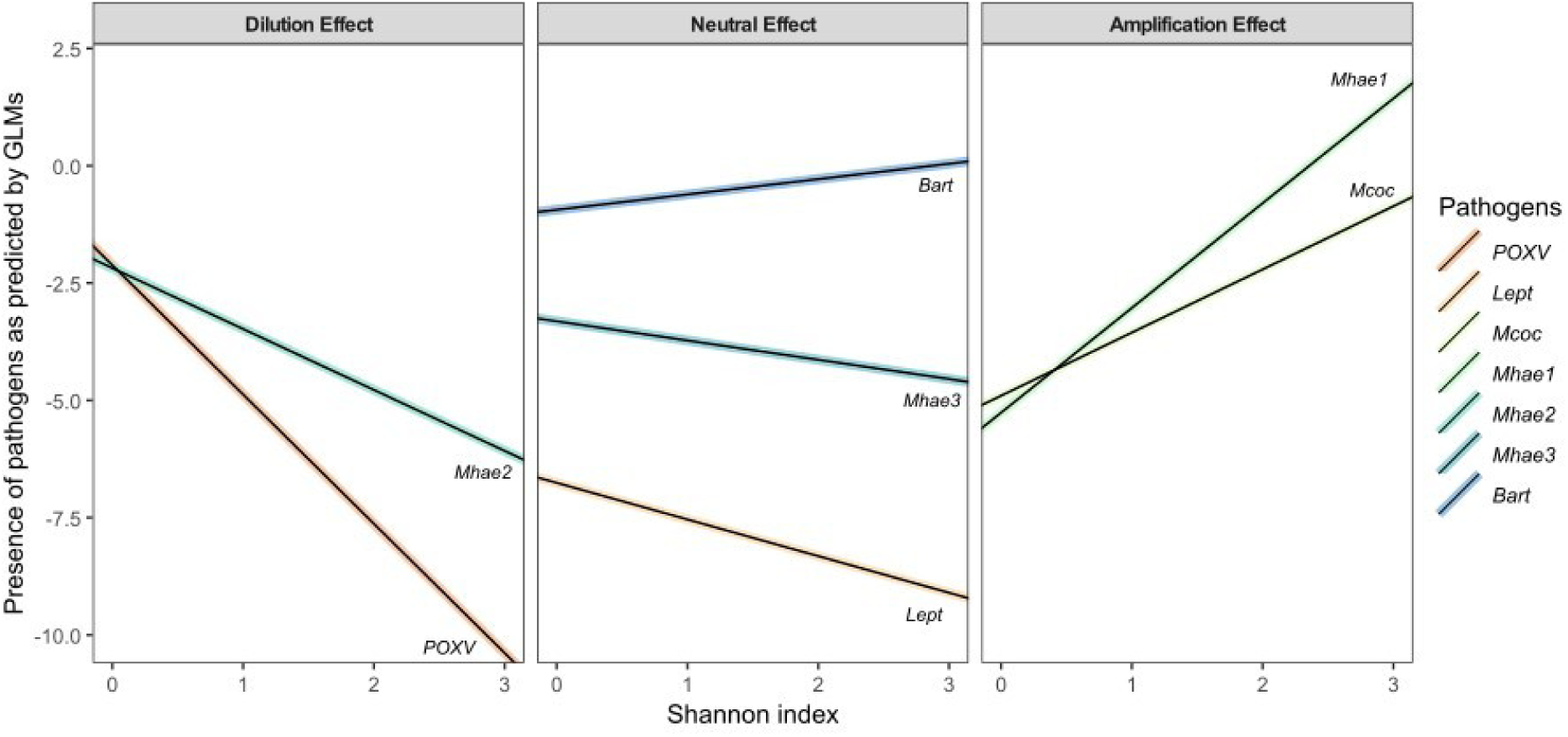
Relationships between pathogen prevalence as predicted by GLMs and host diversity (Shannon index) for each pathogen, with colours corresponding to different pathogens. POXV= antibodies against orthopoxvirus, Lept=*Leptospira* sp., Mcoc= *M. coccoïdes*, Mhae= M. haemomuris (the number following Mhae indicates the corresponding strain). Bart=Bartonella sp.

Conversely, a positive relationship were found between small mammal diversity and the presence of *M. haemomuris* strain 1 (Wald test, β = 2.20, z = 4.45, p = 8.47e-06) *and the presence of M. coccoïdes* (Wald test, β= 1.29, z = 2.91, p = 3.00 e-03) (Fig. 1, Table S1).

Finally, the relationships between the presence of *M. haemomuris* strain 3 (Wald test, β = - 0.14, z = 0.35, p = 0.72), *Bartonella* (Wald test, β = 0.40, zvalue = 1.46, p-value = 0.14) or *Leptospira* (Wald test, β = -0.79, z = -1.63, p = 0.10), and small mammal diversity, were not significant (Fig. 1).

Equivalent relationships were observed when alternative diversity indices or statistical models were used (Diversity indices: Fig. S5, Tables S2 and S3; Correlations: Fig. S6, Table S4; Alternative methods: Table S5).

### Mechanisms underlying the relationship between diversity and pathogen prevalence

To test the hypothesis that a dilution effect occurs when the most competent host species increase in abundance as host community diversity decreases, we implemented a sequential analytical framework including the following steps.

Step 1 - We first assessed interspecific variability in host competence for each pathogen. This was achieved by examining differences in pathogen prevalence across the small mammal species sampled. Our analyses unveiled significant variation in prevalence among host species for each pathogen, indicating substantial differences in host competence within the community (Wald test Chi-square, p = 2.2e-16, Fig. 2, Table S6).

**Fig. 2.**
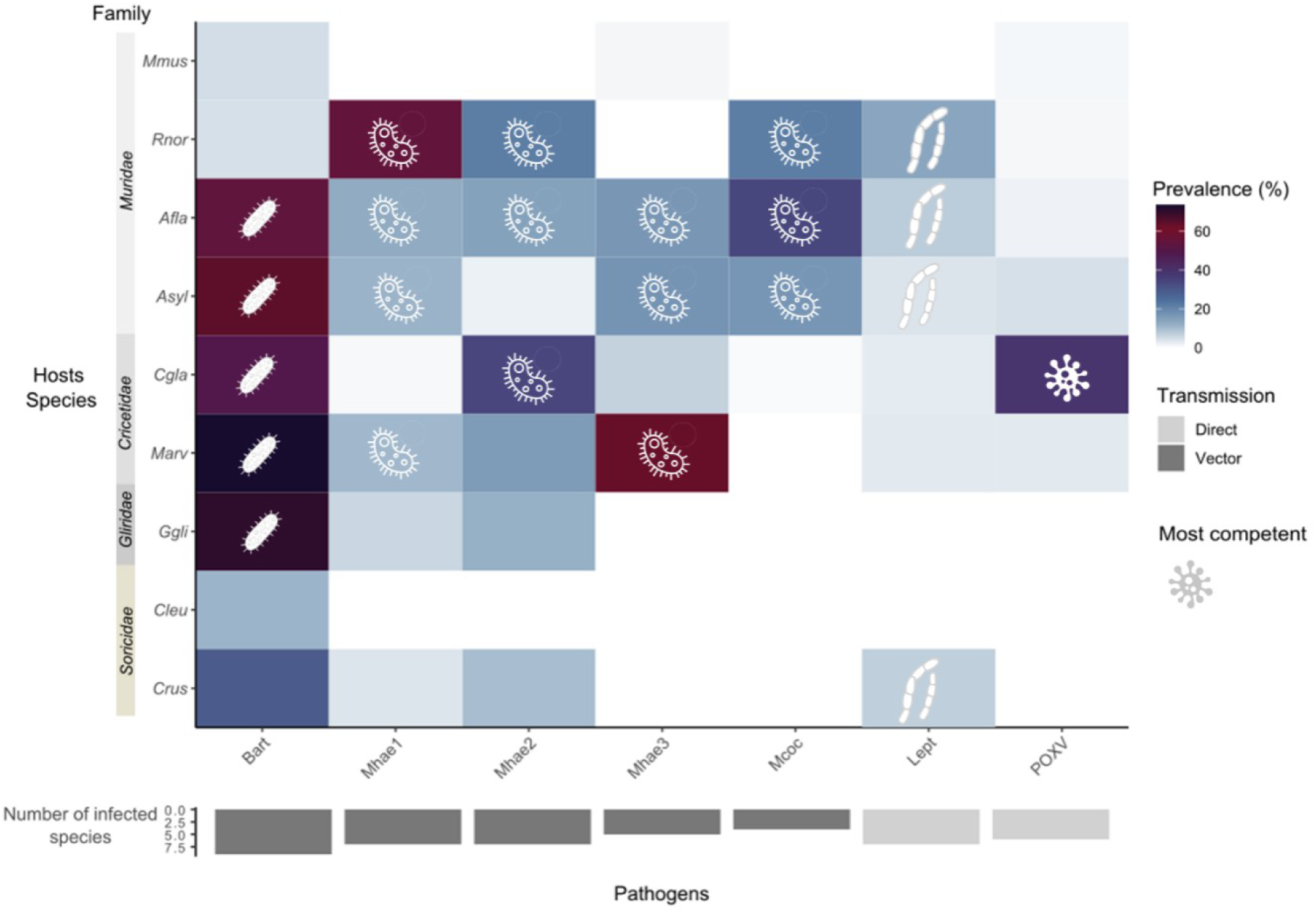
Heatmap showing pathogen prevalence across small mammal host species, with vertical bars indicating family classification. Colour intensity from light blue to dark purple reflects increasing prevalence values. The vertical bar chart displays the number of pathogens detected per host species. The inverted horizontal bar chart indicates the number of host species in which each pathogen was detected. On this chart, light-shaded cells represent directly-transmitted pathogens, whereas dark-shaded ones indicate vector-borne pathogens. Symbols (virus or bacterium icons) indicate the top 10% most competent hosts, based on their average relative prevalence across pathogens. Bart=Bartonella sp., Mhae= M. haemomuris (the number following Mhae indicates the corresponding strain), Mcoc= M. coccoïdes, Lept=Leptospira sp., POXV= antibodies against orthopoxvirus.

Step 2 – We next identified which pathogens exhibited variation in the abundance of their most competent host species as a function of overall host community diversity. We then examined the direction of this relationship.

We considered host species with an average relative prevalence exceeding 10% for a given pathogen as the most competent hosts for this latter (Fig. 2, choice of competence index :Figs. S7 and S8; Table S7). For each pathogen, we then examined how the relative (weighted) abundance of these most competent hosts varied with small mammal community diversity. Our results showed that the abundance of competent hosts was significantly associated with Shannon diversity for all pathogens examined (Fig. 3A, Table S8). Specifically, we observed a decline in the abundance of competent hosts with increasing host diversity for POXV (β = -0.45, t = -6.14, p = 4.29e-06), *M. haemomuris* strain *2* (β = - 0.17, t = -5.80, p = 9.22e-06) and *Bartonella.* The effect size was comparatively smaller for this latter (β = -0.04, t = -3.58, p = 0.001).

**Fig. 3.**
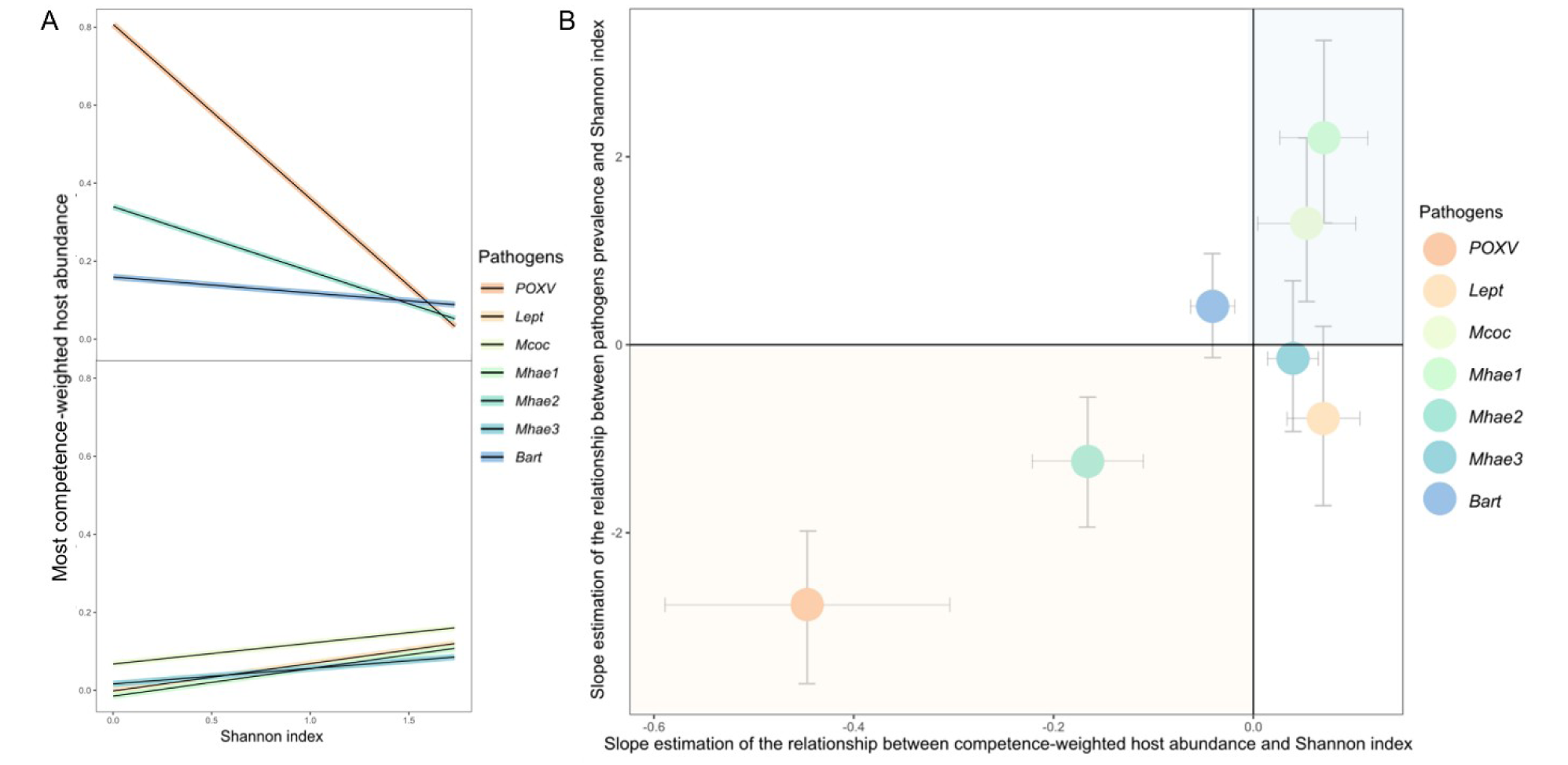
Mechanisms behind the occurrence and strength of dilution effects. **A**. Relationship between the competence-weighted abundance of the most competent host species and small mammal diversity, measured by the Shannon index. **B.** Correlation between the slope of the regression between pathogen prevalence and Shannon index and the estimated slope of the regression between competence-weighted abundance of the most competent species and the Shannon index. Each point represents slope estimates obtained from Generalized Linear Models (GLMs), with error bars indicating 95% confidence intervals derived from the same models. Pathogens are color-coded. The background colour illustrates the direction of the diversity–prevalence relationship: orange indicates a dilution effect (negative slope), blue indicates an amplification effect (positive slope), and white indicates no effect. POXV= antibodies against orthopoxvirus, Lept=*Leptospira* sp., Mcoc= *M. coccoïdes,* Mhae= M. haemomuris (the number following Mhae indicates the corresponding strain), Bart=Bartonella sp.

Conversely, for *M. haemomuris* strain 1, *M. coccoides, M. haemomuris* strain 3, and *Leptospira* sp., the abundance of the most competent hosts increased with greater Shannon community diversity (Fig.3A, Table S8).

Step3 - We investigated whether these observed patterns in competent host abundance could explain the relationship between diversity and pathogen prevalence. We found that the slope of the regression between competence-weighted relative host abundance and small mammal diversity was positively correlated with the slope of the regression between pathogen prevalence and diversity (Fig. 3B). In other words, a dilution effect occurred when the abundance of competent hosts declined with increasing host community diversity, whereas an amplification effect emerged when competent host abundance increased in more diverse communities. Moreover, the steeper the slope of the abundance-diversity relationship, the stronger the corresponding diversity-prevalence relationship (Fig. 3B), suggesting that variation in competent host abundance is a key mechanism driving the direction and strength of diversity–disease associations.

Furthermore, we directly assessed the relationship between the abundance of the most competent hosts and the presence of pathogens for each specific pathogen (Table 1).

**Table 1.** Effects of the most competent host abundance on pathogen presence, accounting for host and environmental factors. Results of the generalized linear models (GLM, binomial distribution) testing the relationship between the abundance of the most competent host species and pathogen presence for each pathogen, while accounting for host-related (age, sex) and environmental (biogeoclimatic conditions, sampling period) confounding factors. Effect estimates are represented using a colour gradient, with more intense shades indicating stronger effects. Yellow highlights statistically significant associations (p < 0.05). POXV= antibodies against orthopoxvirus, Lept=*Leptospira* sp., Mcoc= *M. coccoïdes*, Mhae= M. haemomuris (the number following Mhae indicates the corresponding strain), Bart=Bartonella sp.

| Pathogens | Estimate | z.value | P |
| --- | --- | --- | --- |
| POXV | 4.01 | 8.41 | 2.00e-16 |
| Lept | -12.38 | -2.44 | 1.00e-02 |
| Mcoc | 6.86 | 2.76 | 1.00e-02 |
| Mhae1 | 13.32 | 6.34 | 2.25e-10 |
| Mhae2 | 6.97 | 6.52 | 7.16e-11 |
| Mhae3 | -3.32 | -0.63 | 5.30e-01 |
| Bart | 3.86 | 2.32 | 2.00e-02 |

We demonstrate a positive relationship between the presence of pathogens and the weighted abundance of the most competent hosts (Table 1), regardless of the pathogen, whether associated with dilution or amplification effects. The intensity of this relationship indicates that the abundance of competent hosts plays a critical role in the presence of each pathogen. In contrast, pathogens considered to have no effect in the diversity-pathogen relationship exhibited weak effects, as observed for *Bartonella* sp., or no effect, as seen with *M. haemomuris* strain 3 (Z value= -0.62 and a P =0.52). Notably, *Leptospira* sp. showed a strongly negative slope. Furthermore, the presence of pathogens appeared to be influenced by confounding factors, such as the presence of mature adults (Table S8).

We conducted a comprehensive analysis of both direct and indirect causal relationships using structural equation modeling (SEM; Table S9, Supplementary Note 2) to better understand the complex interactions between host diversity, the abundance of competent hosts, and pathogen prevalence. These analyses significantly supported the patterns previously observed: a negative relationship between host community diversity and the abundance of competent hosts consistently corresponded to a dilution effect, whereas a positive relationship was associated with either an amplification effect or neutral effect. These findings are consistent with the patterns illustrated in Fig. 3B.

## Discussion

This study investigates the various relationships between diversity and pathogen prevalence within a small mammal community sampled across forest sites. Our results reveal a wide range of patterns, with negative associations indicative of a dilution effect, as observed for *Orthopoxvirus* and *Mycoplasma haemomuris* strain 2; positive associations consistent with an amplification effect, for *Mycoplasma coccoïdes* and *Mycoplasma haemomuris* strain 1; and non-significant relationships for *Mycoplasma haemomuris* strain 3*, Leptospira* sp. and *Bartonella*. Altogether, these heterogeneous outcomes indicate that diversity can mitigate or facilitate pathogen transmission, depending on the pathogens investigated in a given context. Such variability has already been reported in previous studies across different host-pathogen systems ^15,27^. However, most findings derive from meta-analyses that aggregate data with different methodologies, spatial scales and environmental conditions. In contrast, our study evaluates these relationships within a single empirical framework over multiple years, in a consistent ecological context, thereby reducing methodological and environmental biases. Importantly, our results were robust across multiple analytical approaches. Similar trends were observed when considering different community-level metrics (species richness and composition). They were also supported by statistical and Structural Equation Modeling, an approach that accounted for both direct and indirect effects.

We did not observe consistent differences in diversity–prevalence relationships based on pathogen transmission modes (direct or vector), taxonomic groups (bacteria, viruses), or host specificity (generalist vs specialist). However, the number of pathogens detected in the study was too low to test these effects statistically. Importantly, we detected variations among different strains of the same pathogen species, *Mycoplasma haemomuris*. These findings underscore the importance of fine-scale taxonomic resolution in pathogen studies and raise concerns about the interpretation of diversity-pathogen prevalence relationships when analyses are conducted at coarse taxonomic levels. This is particularly relevant here for *Bartonella* and *Leptospira* sp., for which our study found no significant diversity effect. Previous work on these small mammal communities have revealed the circulation of multiple *Leptospirosa* species/strains ^28^. Similarly, *Bartonella* studies have shown the possibility of co-infections by multiple species/strains in the same host ^29^. Such taxonomic diversity may mask specific diversity-prevalence associations, thereby explaining the lack of statistical significance in our results for these taxa. Future research should include a large range of pathogens encompassing diverse ecological and epidemiological traits, while ensuring high-resolution taxonomic identification. This approach will be essential for disentangling the complex and context-dependent relationships between diversity and pathogen transmission.

Although competence, defined as an individual’s ability to become infected and transmit a pathogen ^19^, was only indirectly estimated in this study using pathogen prevalence, our approach provides valuable insights. By analyzing average prevalence across seasons and sites for each pathogen and host species, we were able to identify small mammal species that are disproportionally infected by specific pathogens, suggesting interspecific variation in competence. This variation likely reflects specific host–pathogen interactions that could be driven by ecological, physiological or evolutionary traits (Merrill & Johnson, 2020). Understanding competence variability is crucial, as non-competent hosts can influence transmission dynamics, by reducing the abundance or encounter rates of competent hosts ^24^.

Despite competence remains challenging to measure empirically, it remains a key mechanism underlying diversity–disease relationships and thereby should not be overlooked. We argue that incorporating competence, even through proxies such as relative prevalence, is essential in field-based studies. Future efforts could focus on improving its estimation, for example by integrating data on pathogen load. This measure could provide a more accurate estimate of a host’s transmission potential compared to prevalence (Merrill & Johnson, 2020).

Recent meta-analyses have suggested that diversity loss typically leads to an increase in the abundance of highly competent host species – which are generally considered more resistant and tolerant to anthropogenic pressures ^21,25,30^ or widespread generalist hosts that favour transmission opportunities ^31^. However, in our study, we did not observe a consistent positive correlation between host competence and abundance. This may be partly explained by pathogen-specific differences in host competence, as discussed above. Additionally, we hypothesize that the effect of long-term historical anthropogenic pressures have already impacted the small mammal community composition, such that less competent species or more sensitive species may have already been extirpated or are now too rare to be adequately represented in our samples. Consequently, even relatively undisturbed or rural forest communities ^32,33^ are likely dominated by a subset of generalist small mammal species, many of which are at least moderately competent for a range of pathogens. Historical disturbances may limit our ability to detect dilution effects in present-day host communities that have undergone diversity filtering in the past. Future field-based studies could combine trapping and non-invasive approaches to improve the detection of rare and low-abundant host species. This could help reduce sampling biases and provide a more complete understanding of the role of less common hosts on pathogen transmission dynamics.

Our study showed that the strength and direction of the relationship between the abundance of the most competent hosts and small mammal diversity is a strong predictor of the diversity–pathogen prevalence relationship. Specifically, a more pronounced decline in competent host abundance with increasing diversity corresponded to a stronger dilution effect on pathogen prevalence. These findings provide empirical support to theoretical models linking community dynamics (whether additive or substitutive) to dilution and amplification effects ^18^. When the abundance of competent host species decreases with community diversity, transmission risk declines, consistent with dilution effect observed for pathogens such as orthopoxvirus and *Mycoplasma haemomuris* strain 2 ^20,34,35^. Conversely, when both diversity and competent host abundance increase simultaneously, transmission likelihood may increase, leading to amplification effects as seen for *M. haemomuris* strain 1 and *M. coccoides*. This is expected in ecologically unstable or disturbed environments such as urbanized areas ^17,36^.

Local biotic and abiotic conditions may drive shifts in community structure and pathogen ecology, leading to the pathogen-specific responses observed in this study. Such complexity may also result from interspecific competition, which can generate time-lagged or concurrent dilution and amplification effects within the same host community ^17,27^. To disentangle these interacting drivers, we implemented structural equation models (SEM) to evaluate the direct and indirect effects of anthropization and temporal variation on pathogen prevalence. Our results reveal that these factors can influence prevalence both directly (e.g., *Leptospira* sp.) and indirectly, by modulating host diversity and competence. However, we acknowledge that the power of this analysis was constrained by our sampling. Expanding this research across a broader range of ecological contexts is therefore critical to fully understand how environmental and anthropic pressures shape host–pathogen dynamics through their effects on competent host abundance and community diversity ^3^.

In conclusion, this study highlights the importance of maintaining balanced and diverse host communities to limit the prevalence of some pathogens in wild populations and mitigate the associated zoonotic risks. Environments where community composition becomes heavily skewed can experience runaway amplification of disease due to the dominance of highly competent hosts, regardless of overall diversity, whether this imbalance is driven by natural cycles or anthropogenic disturbances. As human activities increasingly disrupt wildlife communities, accelerating non-random species loss, favoring the dominance of some species, and the decline of others, the risk of zoonotic emergence is expected to rise in anthropogenically disturbed environments^37^. Our findings also caution against certain management interventions, such as non-targeted species elimination^38^, which may favor the proliferation of competent hosts and destabilize ecosystems. Instead, we advocate for strategies that preserve both host diversity and stable population abundance, notably through integrated, resilient, and sustainable approaches, such as nature-based solutions and rewilding^39^. As such, a thorough understanding of the ecological relationships among hosts, pathogens and their environment, is crucial before implementing effective zoonosis management ^10^. Future research should further unravel how community composition, host traits, and environmental change interact to shape multi-pathogen dynamics. A greater emphasis on context-dependent effects will be essential for developing robust, adaptive management practices in an era of increasing diversity loss and emerging infectious disease threats.

## Methods

### Small mammal sampling

Small mammal communities were sampled along a forest anthropization gradient to assess how biodiversity influences pathogen prevalence. These species provide an ideal model: their taxonomic diversity, high population density, and ability to occupy a range of habitats (natural and urban^40^) make them both reservoirs of numerous zoonotic pathogens^39^ and ecological bridges between wildlife and humans. Their composition and abundance, sensitive to global change ^40–43^,allow us to study how biodiversity variation along an anthropogenic gradient may affect pathogen circulation.

Sampling was conducted at six sites in eastern France, representing different levels of forest anthropization(Supplementary note 1; Fig. S4: two integral biological reserves (FRFGLA: La Glaciere, FRFGRI: Griffe du Diable); two managed forests (FRFCOR: Cormaranche en Bugey; FRFMIG: Mignovillard); two urban parks (FRPLTO: Lyon, Parc de la Tête d’Or; FRPDLL: Marcy l’Etoile, Domaine Lacroix Laval)).

Captures of small terrestrial mammals were performed during the pre-breeding (spring) and post-breeding (autumn) seasons from spring 2020 to spring 2022 ^44^. Trapping success was estimated for each site and period on the basis of the three first nights of trapping in order to infer the relative abundance of each species captured through the following equation (1) ^45^:

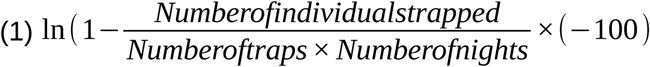

The comprehensive sampling protocol, covering trap setup, species ID, individual dissection and sample storage, are detailed in Pradel et al.,^44^. Detailed morphological information and sexual characteristics were recorded to determine sex and functional age group (immature juvenile versus adult) ^46^.

### Detection of pathogens

#### Detection of antibodies anti-Orthopoxviruses

We applied immunofluorescence assays (IFA) to detect anti-Orthopoxvirus antibodies from heart blood samples (N=1499), using the protocol established by Kallio-Kokko et al. ^47^. Seropositivity was determined by indirect immunofluorescence on Vero E6 cells infected with cowpox virus, with virus-specific antibodies (IgG) detected via fluorescence microscopy.

#### Detection of pathogenic bacteria

DNA was extracted from kidney samples (N = 1500) using BioBasic kits (EZ-10 96 Well Plate Genomic DNA), following the manufacturer’s instructions. Pathogenic Leptospira spp. were detected by real-time PCR (RT-PCR) targeting the lipL32 gene^48^, using a LightCycler LC480 system (Roche Diagnostics) and the protocol described by Dobigny et al. ^49^.

Other potential pathogenic bacteria were detected from spleen samples (N = 1270), an organ involved in blood filtration and immune response. DNA was extracted using the DNeasy 96 Blood and Tissue kit (Qiagen) following the manufacturer’s instructions, and the V4 region of the 16S rRNA gene was amplified and indexed according to Galan et al. ^50^. Samples were then high-throughput sequenced using Illumina MiSeq (2 x 251 bp). Reads were truncated and merged (R1: 180 bp, R2: 120 bp). Sequencing errors were identified and chimeras removed using the dada2 pipeline (Qiime2_2021.11) ^51,52^. Each ASV was assigned to a reference sequence using BLASTN+ ^53^ with the SILVA database (rRNA 138.1) via the FROGS workflow^54^.

Data were filtered following the approach proposed by Galan et al. ^50^ using R v4.2.2 ^55^. Each sample was processed in duplicate with negative and positive controls at each step to monitor contamination. Filters included: (i) removal of ASVs below the maximum counts in negative controls; (ii) elimination of ASVs below the threshold of an "alien" bacterium absent from small mammals, indicating misassignment; (iii) retention of ASVs present in both technical replicates, with sequence counts summed.

We identified pathogenic taxa using the Gideon database (https://www.gideononline.com/) and literature ^56^. Because of 16S sequencing limits, analyses were mostly at the genus level. For *Mycoplasma*, species identification was possible; ASVs matching *Mycoplasma* (syn. *Haemobartonella) haemomuris* and *coccoides* were retained based on Blast (Fig. S5) due to their known pathogenicity and replication in blood and spleen of small mammals. Raw data and scripts are available in the Zenodo repository.

Additional data filtering was applied to ensure sufficient statistical power; only host species represented by more than 10 individuals were retained for subsequent analyses. Furthermore, we focused on pathogens detected across all study sites to minimize biases related to biogeoclimatic effects (Fig. S1). Other pathogens, which are important for surveillance and for understanding key drivers of community structure, are reported in Bouilloud et al.^26^.

### Statistical analyses

#### All analyses described below were performed using R v4.4.0 ^57^

Small mammal diversity was estimated using Shannon index (Tables S6 to S8, Fig. S6) for each site and period (corresponding to a one-year season), which was generated through the vegan package ^58^. To account for environmental factors, we incorporated biogeoclimatic score based on climate data, coordinates, and land use information for each sampling location (Supplementary Note 1).

#### Relationships between pathogen prevalence and small mammal diversity

We tested whether the relationship between pathogen prevalence and small mammal community diversity varied across the different pathogens selected. First, we evaluated the association between the presence of each pathogen and the Shannon index of small mammal communities (other metrics have been analyzed, Tables S3 and S4, Fig. S5) using generalized linear models (GLMs) with a binomial distribution and a logit function. Individual characteristics (functional group and sex) and environmental variables (period and environmental scores for each site) were included in the models. Model selection was performed through an exhaustive exploration (using the MuMIn package). The most parsimonious model(s) were selected with the Akaike information criterion corrected for small sample size (AICc) ^59^ (δAICc < 2). We assessed the goodness-of-fit of each model by analyzing the deviation and dispersion of model residuals from normality using the DHARMa package ^60^ and checked the variance inflation factor (VIF) to address multicollinearity.

#### Analytical framework for investigating the relationships between diversity-prevalence mechanisms

To investigate the underlying mechanisms of the dilution effect, we tested the hypothesis that, under such scenarios, the most competent host species tend to become more abundant as host community diversity declines. To do so, our approach comprised the following three steps.

Step 1: We a*ssessed variation in host species competence across pathogens and quantifying host competence*. We applied an optimized generalized linear mixed model (GLMM) with pathogens presence/absence as the response variable, pathogen taxa, host species, environmental scores, sampling period and individual factors (functional group and sex) as fixed effects, and individual identity as a random effect. To assess the variation in host species competence for each pathogen, we investigated the propensity of pathogens to preferentially infect certain host species and whether this tendency varied depending on the pathogen species. This was achieved by modeling an interaction between pathogen species and host species, followed by a post-hoc test to more precisely compare host species exhibiting different levels of “competence” for a specific pathogen.

To quantify host species’ competence for a given pathogen, we used a proxy that based on infection prevalence to compare this latter among species. Specifically, we calculated the mean relative prevalence of infected host species across all sampling sites and periods (N=23) for the pathogen of interest. After initial threshold analyses and comparison with the more competent host species identified by GLMM post-hoc results (Supplementary Note1, Fig. S7, Table S6), we classified host species with an average relative prevalence greater than 10% as the most "competent" or preferentially infected hosts (see alternative thresholds in Table S7 and Fig. S8).

*Step 2:* We examined the relationship between the competent host abundance and the overall host community diversity. For each pathogen, we fitted generalized linear models (GLMs) with a Gaussian error distribution, using a simple linear regression framework without additional covariates to preserve statistical power given the limited number of site-period combinations (N = 23).

The explanatory variable was host community diversity, measured using the Shannon diversity index. The response variable was defined as the cumulative relative abundance of host species identified as most competent for each pathogen (i.e., those with a mean relative prevalence >10%), weighted by their average infection prevalence across all sites and periods. This weighting provided a proxy for host competence intensity.

To ensure the robustness of the results, we also tested alternative metrics of competent host abundance (Table S8).

Step 3: We analysed the relationships between competent host abundance—both direct and mediated by community diversity—and pathogen presence. Due to the limited number of pathogens (N = 7, corresponding to 7 points, Fig. 3), formal statistical testing of whether the relationship between the abundance of the most competent host species and species diversity (Shannon index) explains the observed association between host diversity and pathogen prevalence was constrained. We therefore applied two complementary modelling approaches.

First, we used a generalized linear model (GLM, binomial distribution) to evaluate the direct effect of competent host abundance on the pathogen presence, separately for each pathogen. Models accounted for potential confounding factors, including host characteristics (age, sex) and environmental variables (biogeoclimatic context and sampling period), and followed the same framework previously used for variable selection and model fitting. The rationale is that if both relationships A ∼ B and C ∼ B are supported—where A is the abundance of competent hosts, B is species diversity, and C is pathogen presence—then a significant positive A ∼ C association provides indirect evidence for a mediation mechanism.

Second, we fitted structural equation models (SEMs) incorporating environmental covariates to simultaneously estimate the direct and indirect effects of species diversity and competent host abundance on pathogen presence, separately for each pathogen (Fig. S9). This approach is particularly suited for testing complex causal hypotheses, as it allows for the detection of mediation pathways that cannot be captured by simpler models (such as in (i)), and facilitates the identification of potential dilution or amplification mechanisms (see details in Supplementary note 2, Table S9, Fig. S9).

## Supporting information

Description of Addtional Supplementary Files

## Data availability

The raw data have been deposited on Zenodo and are available at https://zenodo.org/records/12518286.

## Code availability

R scripts have been deposited on Zenodo and are available at 10.5281/zenodo.15944869.

## Acknowledgements

Thanakorn Niamsap, Sofia Greilich, Ella Ahviainen, Ella Lintunen, Akseli Valta, Hayder Assad, Eetu Sironen, Vinaya Venkat provided assistance with the IFA. Their valuable contributions and warm hospitality in their laboratory are greatly appreciated.

## Authors contributions

MB: Conceptualization; Investigation; Data curation; Formal analysis; Methodology; Visualization; Writing—original draft. MG: Conceptualization; Methodology; Investigation; Supervision; Data curation; Formal analysis; Writing—review & editing. JP: Data curation; Resources; VS: Conceptualization; Investigation; review & editing; NC: Project administration; Conceptualization; Funding acquisition; Methodology; Supervision; Validation; Writing—review & editing. BR: Conceptualization; Funding acquisition; Methodology; Supervision; Validation; Writing—review & editing.

## Supplementary information

Description of Addtional Supplementary Files, File: *Summary_FigTabNotes_Bouilloud.pdf*

Supplementary Figures, File: *SupplementaryFigures_Bouilloud.docx*

Supplementary Tables, File: *SupplementaryTables_Bouilloud.xlsx*

Supplementary Note 1, File: SupplementaryNote1_Bouilloud.docx

Supplementary Note 2, File: SupplementaryNote2_Bouilloud.docx

