## Supplementary Note 2 for "Host competence–abundance relationships drive the dilution effect across multiple small mammal-borne pathogens"

### Supplementary note 2: Materials and Results of Structural Equation Modeling

Methods

Understanding diversity–prevalence relationships: direct and indirect effects of environment and competent host abundance

We employed a confirmatory path analysis using the PiecewiseSEM package^1^ to explore the interactions between the environment (gradient of forest anthropization), host community biodiversity, and the presence/absence of pathogens. The structural equation modelling (SEM) allowed us to examine the direct effects of the environment (as predictor variables) on pathogens’ presence (response variable), as well as their indirect effects through host biodiversity changes (responses variables: Shannon diversity or relative abundance of most competent hosts), which in turn (biodiversity variables as predictor variables) influenced pathogens’ presence (response variable) (Fig. S9). The path analysis involved fitting three generalized linear models (GLMs) with different response distributions (binomial for presence/absence of pathogens, Gaussian for biodiversity metrics). To enhance the performance of the SEM, we conducted individual tests for each path to identify the best-fitting GLMs. These optimal models were then integrated into the global SEM. The fit of this global SEM was assessed using Fisher's C-test (p > 0.05).

Results

The best SEM obtained for POXV (Fischer’s C = 0.35, p-value = 0.84, R²presence = 0.21) and Mhae2 (Fischer’s C = 5.87, p-value = 0.44, R²presence = 0.06) showed that a decrease in biodiversity led to an increase in pathogen presence (POXV: Std. Estimate = -0.15, p-value = 1.00e-03, Mhea2: Std. Estimate = -0.17, p-value <1.00e-03), as expected under the dilution effect. This decrease in biodiversity was associated with a higher relative abundance of the most competent hosts (POXV: Std. Estimate = - 1.16, p-value < 1.00e-03, Mhea2: Std. Estimate = -1.10, p-value < 1.00e-03). These models also revealed that anthropization and other environmental factors had a significant influence on the presence of pathogens through either direct or indirect interactions via their impact on the abundance of the most competent hosts (Table S9).

The best models provided a good fit for explaining the presence of *Mhea1* (Fischer’s C= 13.17, p-value=0.106, R²=0.07) and *Mcoc* (Fischer’s C= 4.43, p-value=0.35, R²=0.02). In all cases, we did not observe a direct effect of diversity on pathogen presence (Table S9), but an indirect effect through the abundance of most competent hosts. This abundance was positively correlated with biodiversity (Mhea1: Std. Estimate = 0.08, p-value < 1.10-3, Mcoc: Std. Estimate = 0.13, p-value = < 1.10-3) and pathogen presence (Mhea1: Std. Estimate = 0.29, p-value < 1.10-3, Mcoc: Std. Estimate = 0.19, p-value = < 1.10-3) (Table S9). Other environmental factors could also influence pathogen presence through the abundance of competent hosts or diversity but not directly (Table S9).

For *Bartonella,* the best SEM (Fisher's C = 7.29, p-value = 0.70, R²presence=0.02) revealed that biodiversity positively influenced the presence of this bacteria (Std. Estimate = 0.15, p-value = < e-03), but the abundance of the most competent hosts was negatively correlated with biodiversity and anthropization (Table S9). Other environmental factors could also influence *Bartonella*’s presence, but only indirectly through their impact on the abundance of the most competent hosts or on biodiversity (Table S9).

For Mhea3, the best SEM (Fisher's C = 3.84, p-value = 0.43, R²presence=0.03) showed that biodiversity had a negative influence on pathogen presence and was positively correlated with the abundance of the most competent hosts (Std. Estimate =- 0.21, p-value = < e-03). Anthropization negatively influenced Mhea3 presence, and other environmental factors had both direct and indirect positive effects on this pathogen (Table S9).

The best SEM for *Leptospira* (Fischer’s C= 1.33, p-value= 0.514, R²presence=0.11) showed that the presence of this pathogen was directly influenced by the sampling period and environmental effects, but not by small mammal biodiversity or abundance of the most competent hosts (Table S9).
