## Supplementary Note 1 for "Host competence–abundance relationships drive the dilution effect across multiple small mammal-borne pathogens"

### Supplementary note 1 : Detailed methodological information

Site characterization based on biogeoclimatic and anthropization gradients

To integrate environmental factors into our analyses, we defined two scores—biogeoclimatic and anthropogenic—using Principal Component Analysis (PCA) with the FactoMineR package ^1^, Biogeoclimatic data were obtained by combining geographical coordinates, land use information from the Corine Land Cover map (CORINE Land Cover, 100 m resolution), and bioclimatic indices extracted from the CHELSA database (1 km resolution, <https://chelsa-climate.org/>) ^2^ for each site, using the coordinates of the trap line's barycenter.

To quantify the anthropization gradient, we used the Corine Land Cover map and estimated forest fragmentation (Corine Land Cover for forests, 10 m resolution) with the Landscapemetric package^3^ . Additionally, we considered the level of forest exploitation by classifying forests into different management practices, ranging from no management (level 0) to various degrees of intervention for economic, social, or infrastructural purposes (levels 1 to 5).

Characteristics of small mammal diversity metrics and community composition indices

Different diversity indices of small mammals were developed to ensure the robustness of the results, particularly in analyses exploring the relationship between host diversity and pathogen prevalence. The specific richness and Shannon index were estimated using the vegan package. The composition of small mammal communities was evaluated through a Bray-Curtis dissimilarity matrix generated with vegan package, and it was subsequently ordered using a Principal Coordinate Analysis (PCoA) done with the stats package. For each site and sampling period, the score along the first axis was retained as an index of small mammal community composition.

Correlation analyses between small mammal diversity metrics (species richness, Shannon index, and community composition score) and the anthropization and biogeoclimatic scores described above were conducted using Pearson’s correlation test with the cor_test function from the rstatix package^4^ . These analyses revealed significant correlations among several indices, notably between biogeoclimatic and anthropization scores, justifying the inclusion of only one biogeoclimatic factor in certain models due to collinearity. Additionally, significant correlations between Shannon diversity, species richness, and community composition score were observed; however, Shannon diversity was selected as the main diversity metric in subsequent analyses due to its strong and consistent associations. These supplementary results confirm the overall pattern identified in our main analyses.

Statistical analyses of the relationship between host diversity and pathogen prevalence

We examined whether our results were robust to the indices chosen and the model selected. We explored various calculation methods, as well as different levels and types of diversity indices.

We first evaluated the relationship between pathogen presence and small mammal diversity (species richness or community composition score), using generalized linear models (GLMs) with a binomial distribution and a logit link function, for each pathogen. Models included individual-level covariates (functional group, sex) and environmental variables (sampling period and biogeoclimatic scores). Model selection was based on AICc (ΔAICc < 2), and model assumptions were verified through residual diagnostics and multicollinearity checks.

We adopted a unified approach by using a single model to examine the link between diversity (Shannon) and pathogen presence, incorporating all pathogens (i.e., ANCOVA). We applied a generalized linear mixed model (GLMM) with individuals as random effects, alongside fixed environmental and individual factors. The evaluation of goodness-of-fit and model selection methods were consistent. We performed a post-hoc test using emmeans_test with rstatix package^4^ to compare pairs of slopes between pathogens. Benjamini-Hochberg corrections for multiple tests were used to assess the significance of interactions. Our findings indicated comparable trends, but the effect was attenuated due to the distinct impact of each pathogen on infection numbers.
