## Supplementary Figures for "Host competence–abundance relationships drive the dilution effect across multiple small mammal-borne pathogens"

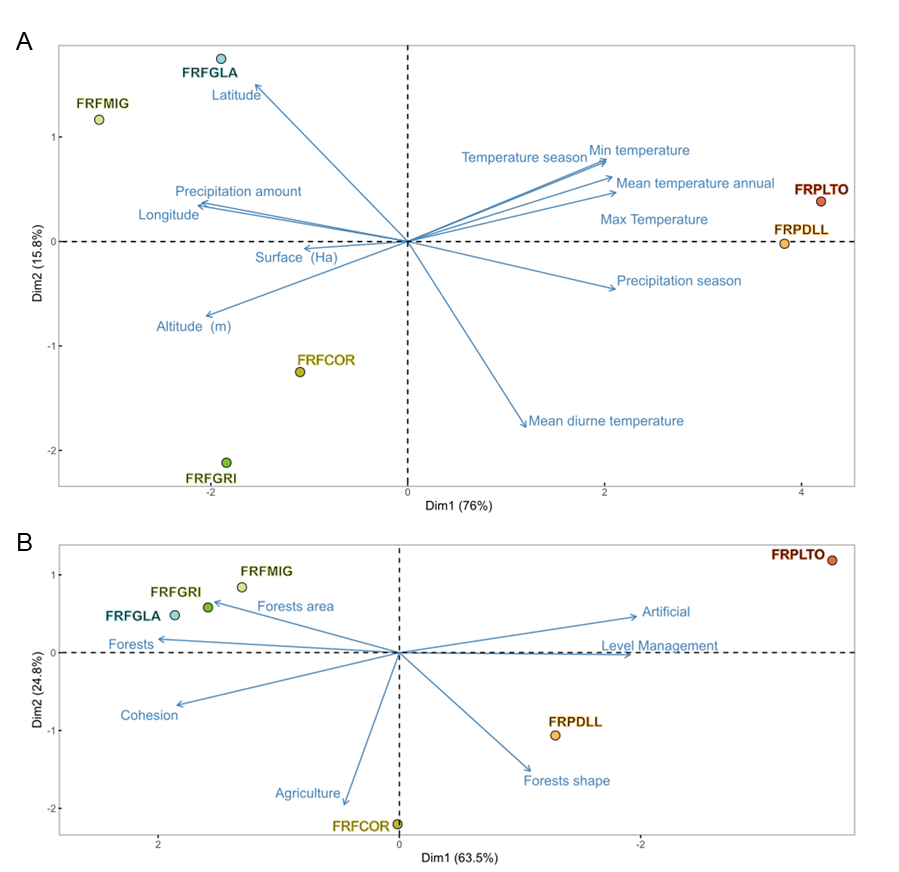

**Fig. S1. Principal Component Analysis of biogeoclimatic and anthropogenic factors across study sites.** A. Principal Component Analysis (PCA) based on biogeoclimatic characteristics. Various factors are derived from GPS coordinates and the Chelsa database (definitions on this site <https://chelsa-climate.org/bioclim/>); B. PCA based on anthropogenic factors. FRPLTO: Lyon, Parc de la Tête d'Or (Rhône); FRPDLL: Marcy l'étoile, Domaine Lacroix Laval (Rhône); FRFCOR: Cormaranche en Bugey (Ain); FRFGRI: Arvière, La Griffe au diable (Ain); FRFMIG: Mignovillard (Jura); FRFGLA: Esserval-Tartre, La Glacière (Jura).

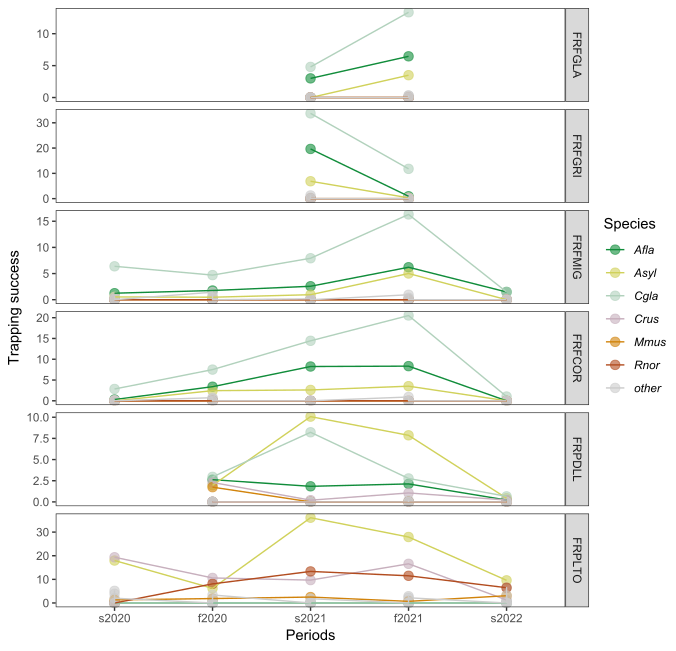

**Fig. S2.** **Average trapping success rate observed for the most abundant species across the six study sites over time.** Species are represented by colors (Asyl = Apodemus sylvaticus; Afla = Apodemus flavicollis, Cgla = Clethrionomys (syn. Myodes) glareolus, Crus = Crocidura russula, Mmus = Mus musculus, Rnor = R norvegicus). Periods are categorized by season (s = spring and f = fall). FRPLTO: Lyon, Parc de la Tête d'Or (Rhône); FRPDLL: Marcy l'étoile, Domaine Lacroix Laval (Rhône); FRFCOR: Cormaranche en Bugey (Ain); FRFGRI: Arvière, La Griffe au diable (Ain); FRFMIG: Mignovillard (Jura); FRFGLA: Esserval-Tartre, La Glacière (Jura).

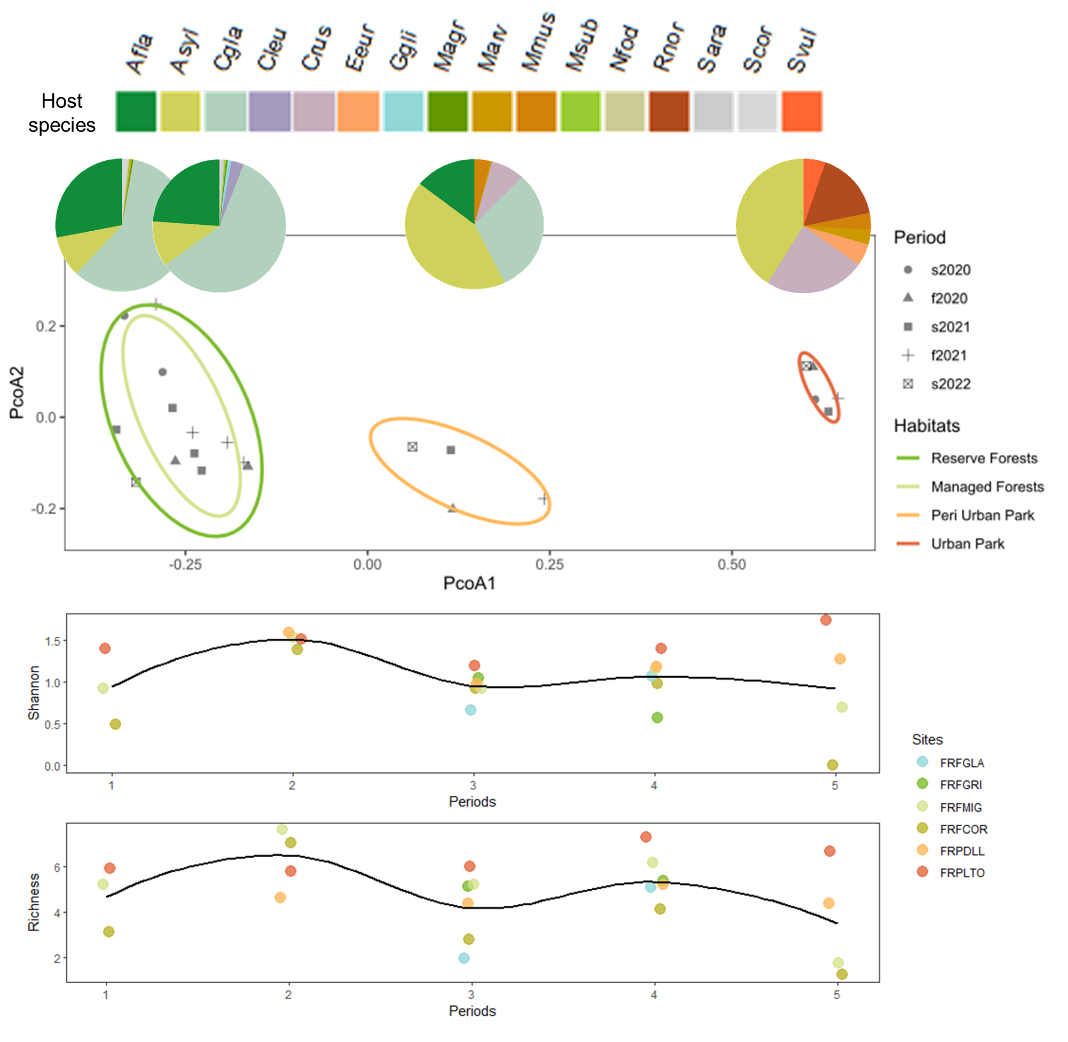

**Fig S3. Variation in small mammal community composition and diversity across habitats and time. A.** Principal Coordinate Analysis (PCoA) was used to visualize the variation in community composition of the small mammal communities sampled at various sites and periods. Each point on the plot represents a community (site*period), with different shapes indicating the sampling period, arranged in order, with 1 representing spring 2020 and 5 representing autumn 2022. Ellipses are used to highlight different habitat types: dark green for Reserve forests (FRFGRI & FRFGLA), light green for Managed forests (FRFCOR & FRFMIG), orange for peri-urban parks (FRPDLL), and red for Urban parks (FRPLTO). Within each ellipse, we presented the mean relative abundance of small mammal species (represented by different colors) with pie charts.

The PCOA revealed that the primary difference in community composition was driven by habitat type (Axis 1). While no significant difference was observed between reserve and managed forest sites, changes in community composition were found between rural forests, the peri-urban (FRPDLL) and the urban (FRPLTO) parks. This axis was significantly correlated with the anthropisation score (correlation = 0.95, p-value = 7.50e-12). In a lesser extent, we also observed variations in small mammal community composition through time.

**B.** The Shannon index and species richness are plotted across periods and sites, with the colors of the points corresponding to different sites. The lines represent the overall temporal trends.

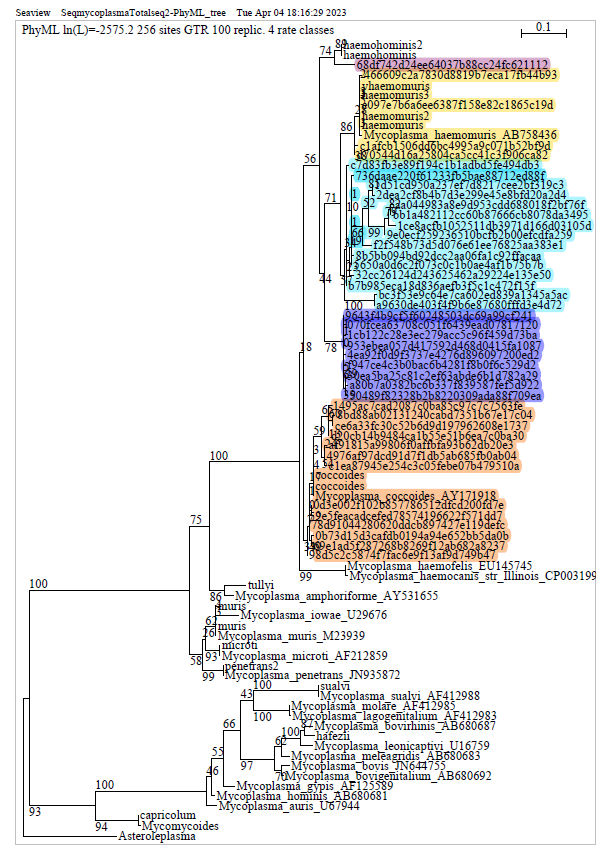

**Fig. S4. Phylogenetic tree of Mycoplasma ASVs circulating in small mammal blood. The 16S V4 sequences obtained in this study were aligned with reference sequences retrieved from NCBI (fasta files, labeled as** Mycoplasma_species_CodeID**, e.g., M23939) using MUSCLE in SeaView^7^. A maximum likelihood phylogenetic tree was inferred in SeaView with PhyML under the GTR model with four rate categories (GTR + 4Γ) and 100 bootstrap replicates. The tree was rooted using pre-selected outgroup sequences from other** Mycoplasma **species. The log-likelihood of the inferred tree was lnL = 2575.2 over 256 aligned sites. Each ASV from our metabarcoding dataset is labeled by its ASV number, with colors indicating distinct** Mycoplasma haemomuris **strains.**

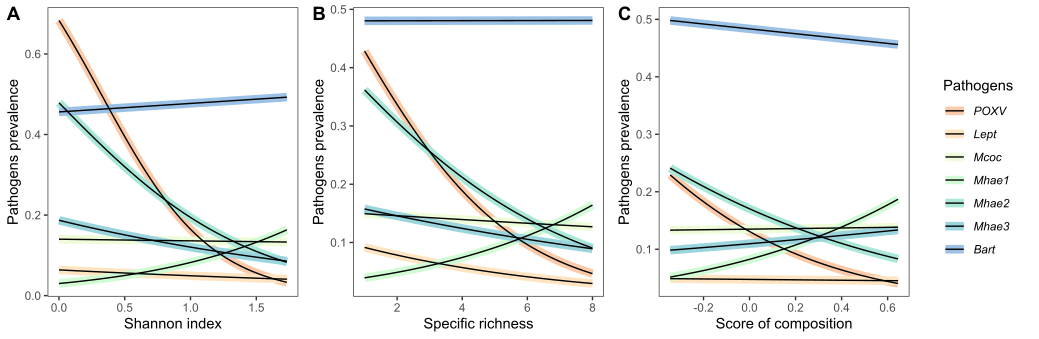

**Fig.S5. Relationships between pathogen prevalence and different indices of host diversity for each pathogen, with colors representing different pathogens.** A. Shannon index, B. Richness, C. Community composition score. Pathogens tested were Or*thopoxvirus (POXV)*, *Leptospira spp. (Lept),* *Bartonella spp. (Bart)*, *Mycoplasma coccoides* (*Mcoc*) and *Mycoplasma haemomuris strains 1–3 (Mhae1–Mhae3)*.

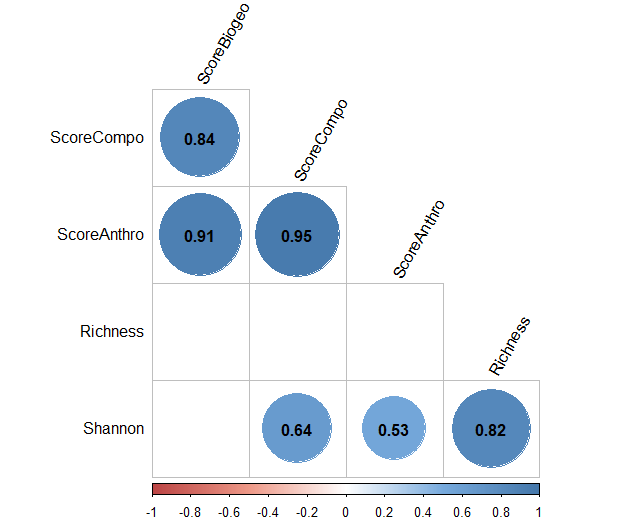

**Fig. S6. Correlation plot between small mammal diversity and environmental scores.** This plot shows correlations between diversity indices—including species richness, Shannon index, and community composition scores—and environmental scores, which comprise both biogeoclimatic variables and anthropogenic factors. Correlations were assessed using Pearson correlation with the cor_test function from the **rstatix** R package^1^ .

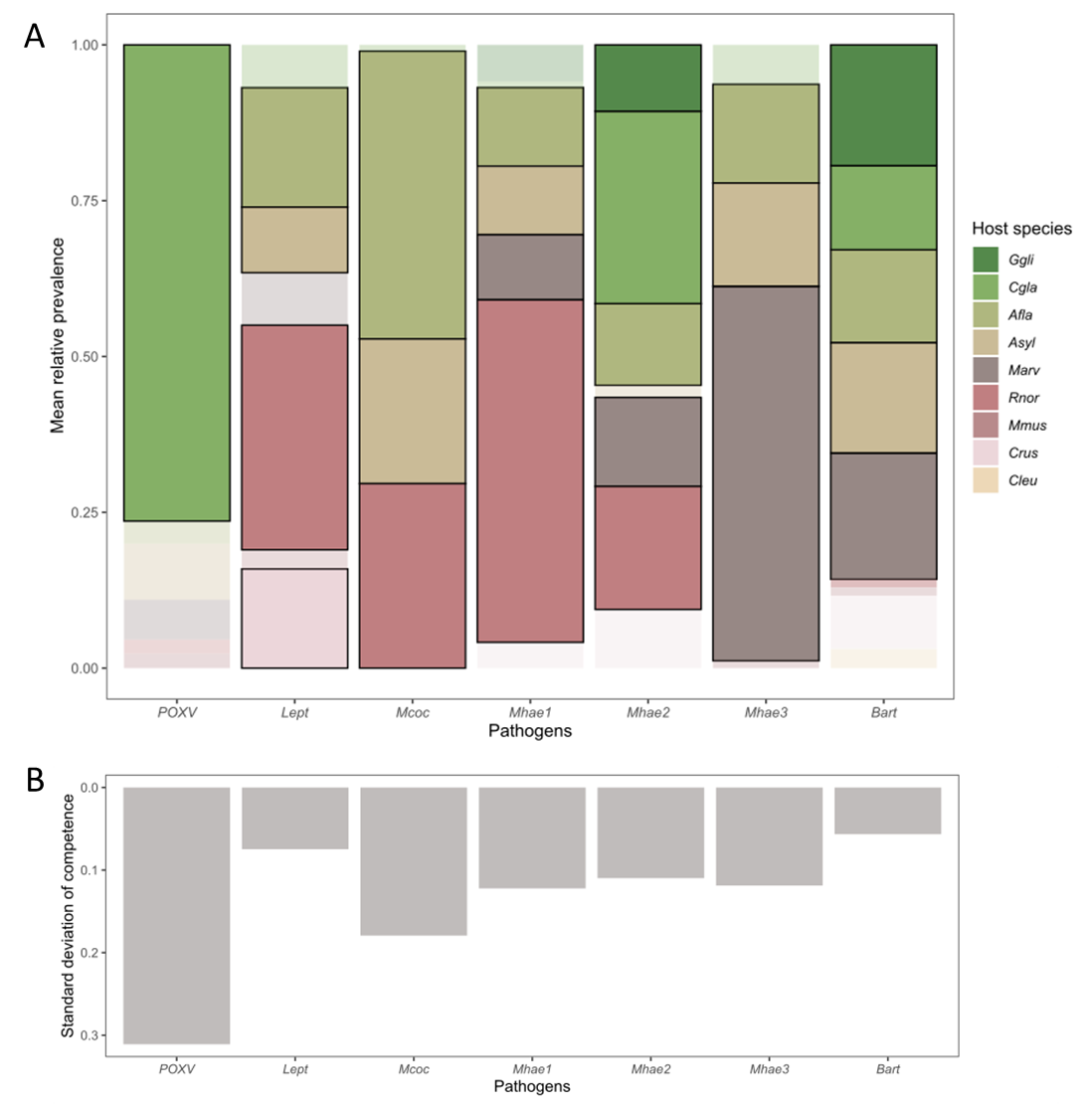

**Fig.S7. Host-specific prevalence and competence across pathogens in small mammals.** Barplot of A. the mean relative prevalence of host species for each pathogen, with colors indicating the host species, and a black outline indicating host species considered as the most competent hosts with a mean prevalence >10% of the total prevalence for that pathogen. Colors of other species are faded. B. Standard deviation of host competence for each pathogen. Pathogens tested were Or*thopoxvirus (POXV)*, *Leptospira spp. (Lept),* *Bartonella spp. (Bart)*, *Mycoplasma coccoides* (*Mcoc*) and *Mycoplasma haemomuris strains 1–3 (Mhae1–Mhae3)*. The small mammal species included in this study were *Glis glis (Ggli), Clethrionomys glareolus (Cgla), Apodemus flavicollis (Afla), Apodemus sylvaticus (Asyl), Microtus arvalis (Marv), Rattus norvegicus (Rnor), Mus musculus (Mmus), Crocidura russula (Crus), and Ctenophorus leucogaster (Cleu).*

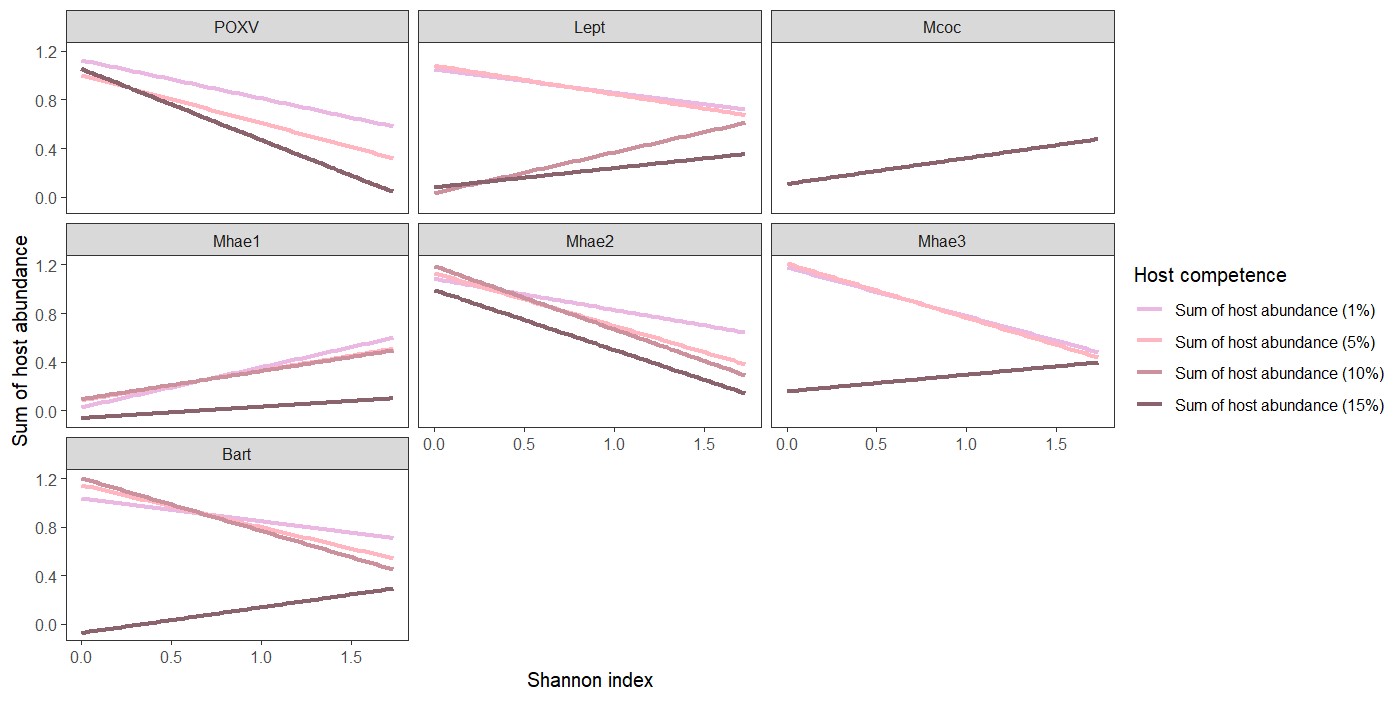

**Fig. S8. Effect of the mean relative prevalence (MRP) threshold defining the most competent hosts on the relationship between their abundance and Shannon diversity.** Abundances were summed for species exceeding each threshold (see **Supplementary Table 7**). The pink gradient indicates average pathogen prevalence from 1 % (pale) to 15 % (dark). Trends are largely consistent across thresholds, except for Bartonella (Bart), Leptospira (Lept), and Mycoplasma haemomuris strain 3 (Mhae3), where only the 15 % threshold for Bart and Mhae3, and the 10 % threshold for Lept, deviate, consistent with GLMM results.

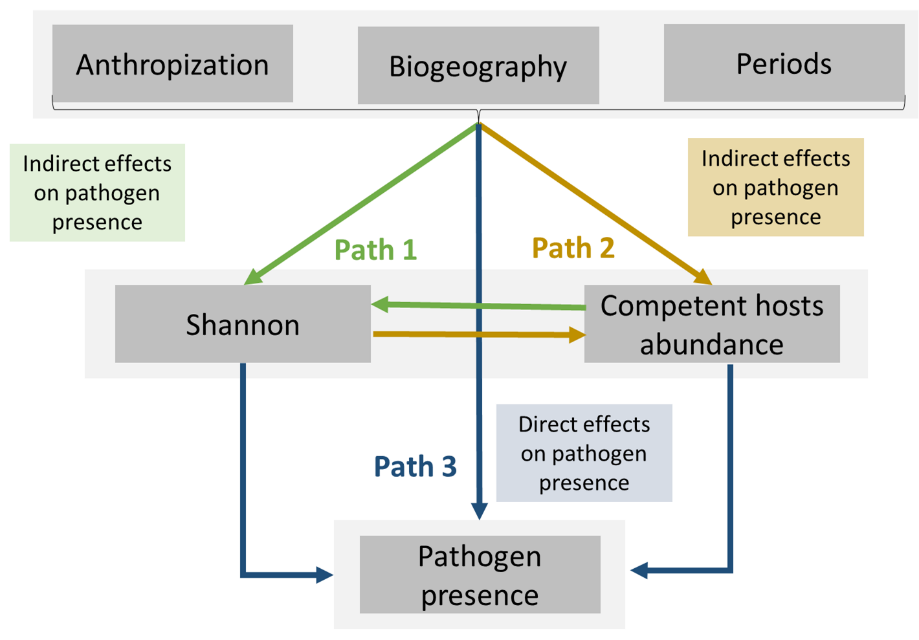
**Fig. S9. Path analysis of direct and indirect effects on pathogen presence.** Arrows and colors indicate the type of effect on pathogen presence. Blue arrows represent direct effects, yellow arrows represent indirect effects mediated by the abundance of the most competent host, and green arrows represent indirect effects mediated by the Shannon diversity index.

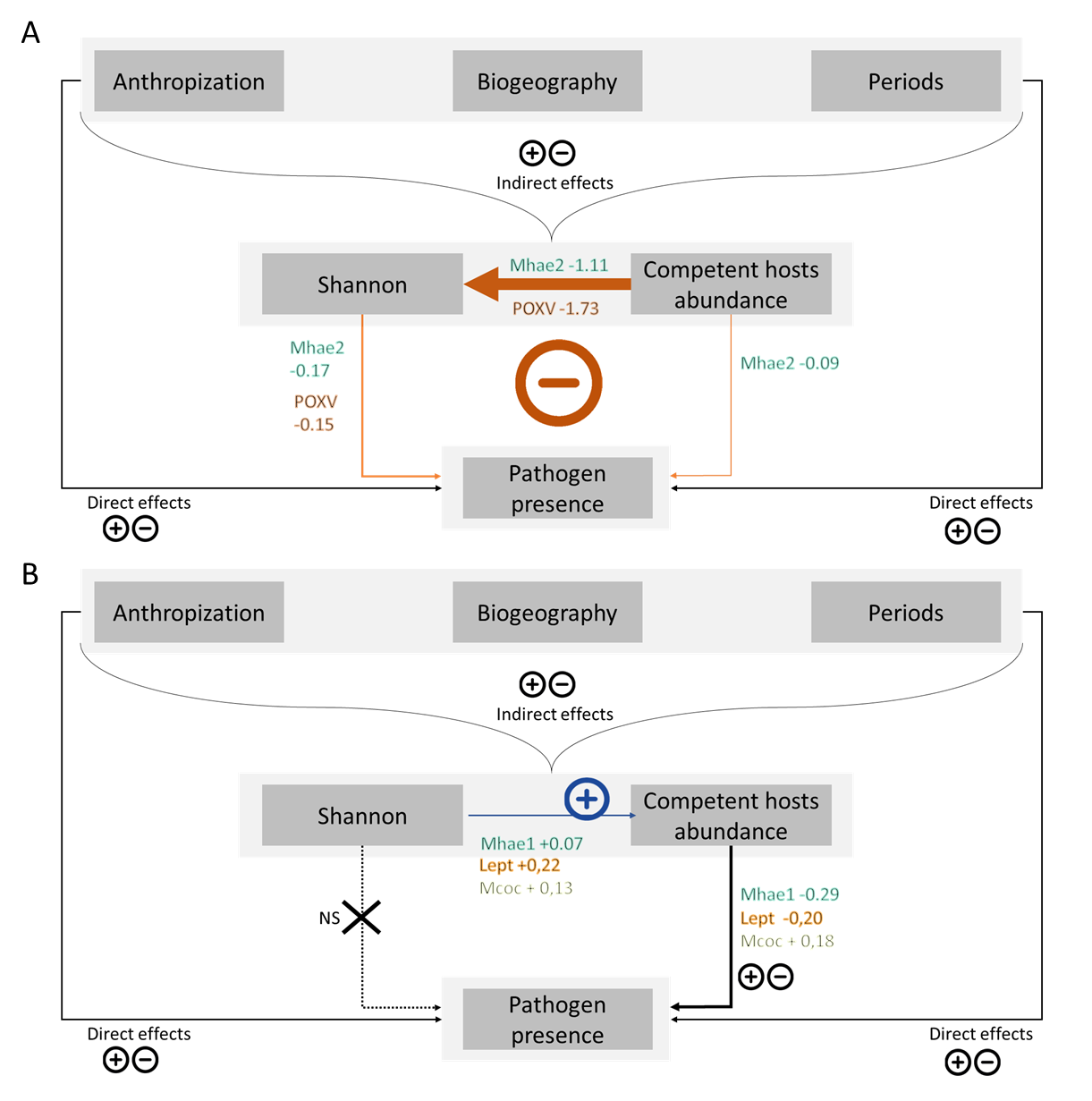
**Fig. S10. Summary of structural equation models (SEMs) illustrating factors shaping pathogen presence.** SPanel A shows dilution effects for Orthopoxvirus (POXV) and Mycoplasma haemomuris strain 2 (Mhae2), while Panel B shows amplification effects for Mhae2 and Mycoplasma coccoides (Mcoc). Arrows indicate relationships: orange for negative, blue for positive, and black for relationships that could be either. Crosses and dotted arrows denote non-significant effects. See Table S9 for details of the SEMs.
