## Supplementary material for "Host competence–abundance relationships drive the dilution effect across multiple small mammal-borne pathogens": Description of Addtional Supplementary Files

### Summary of Tables, Figures, and Notes – Bouilloud et al.

#### Main Text Figures and Table

- **Fig. 1.** Relationships between pathogen prevalence predicted by GLMs and host diversity (Shannon index) for each pathogen.
- **Fig. 2.** Heatmap showing pathogen prevalence across small mammal host species. Vertical bars indicate family classification.
- **Fig. 3.** Mechanisms underlying the occurrence and strength of dilution effects.
- **Table 1.** Effects of the most competent host abundance on pathogen presence, accounting for host and environmental factors (GLMs, binomial distribution).

**File:** *MainText_FigTable_Bouilloud.docx*

#### Supplementary Tables

- **Table S1.** GLM results – effect of host Shannon diversity and confounding factors on pathogen presence.
- **Table S2.** Pearson correlation tests between small mammal community diversity indices and environmental factors.
- **Table S3.** GLM results – effect of small mammal richness and confounding factors on pathogen presence.
- **Table S4.** GLM results – effect of small mammal community composition score and confounding factors on pathogen presence.
- **Table S5.** GLMM results – relationships between pathogen presence and small mammal community structure estimated from different metrics.
- **Table S6.** GLMM results – relationships between pathogen presence and host species.
- **Table S7.** Host competence classification based on mean relative prevalence (MRP) and GLMM significance.
- **Table S8.** GLM results – effects of abundance of the most competent hosts on small mammal community diversity and pathogen prevalence.
- **Table S9.** Best-fit SEM results for pathogen prevalence in small mammal hosts.

**File:** *SupplementaryTables_Bouilloud.xlsx*

### Supplementary Figures

- **Fig. S1.** Principal Component Analysis of biogeoclimatic and anthropogenic factors across study sites.
- **Fig. S2.** Average trapping success rate observed for the most abundant species across the six study sites over time.
- **Fig. S3.** Variation in small mammal community composition and diversity across habitats and time.
- **Fig. S4.** Phylogenetic tree of *Mycoplasma* ASVs circulating in small mammal blood.
- **Fig. S5.** Relationships between pathogen prevalence and different indices of host diversity for each pathogen, with colors representing different pathogens.
- **Fig. S6.** Correlation plot between small mammal diversity and environmental scores.
- **Fig. S7.** Host-specific prevalence and competence across pathogens in small mammals.
- **Fig. S8.** Effect of the mean relative prevalence (MRP) threshold defining the most competent hosts on the relationship between their abundance and Shannon diversity.
- **Fig. S9.** Path analysis of direct and indirect effects on pathogen presence.
- **Fig. S10.** Summary of structural equation models (SEMs) illustrating factors shaping pathogen presence.

**File:** *SupplementaryFigures_Bouilloud.docx*

### Supplementary Notes

- **Supplementary Note 1:** Detailed methodological information
  **File:** *SupplementaryNote1_Bouilloud.docx*
- **Supplementary Note 2:** Materials and results of structural equation modeling
  **File:** *SupplementaryNote2_Bouilloud.docx*
